# Multiplexed detection of SARS-CoV-2 genomic and subgenomic RNA using *in situ* hybridization

**DOI:** 10.1101/2021.08.11.455959

**Authors:** Kofi K. Acheampong, Dylan L. Schaff, Benjamin L. Emert, Jonathan Lake, Sam Reffsin, Emily K. Shea, Courtney E. Comar, Leslie A. Litzky, Nigar A. Khurram, Rebecca L. Linn, Michael Feldman, Susan R. Weiss, Kathleen T. Montone, Sara Cherry, Sydney M. Shaffer

## Abstract

The widespread Coronavirus Disease 2019 (COVID-19) is caused by infection with the novel coronavirus SARS-CoV-2. Currently, we have a limited toolset available for visualizing SARS-CoV-2 in cells and tissues, particularly in tissues from patients who died from COVID-19. Generally, single-molecule RNA FISH techniques have shown mixed results in formalin fixed paraffin embedded tissues such as those preserved from human autopsies. Here, we present a platform for preparing autopsy tissue for visualizing SARS-CoV-2 RNA using RNA FISH with amplification by hybridization chain reaction (HCR). We developed probe sets that target different regions of SARS-CoV-2 (including ORF1a and N) as well as probe sets that specifically target SARS-CoV-2 subgenomic mRNAs. We validated these probe sets in cell culture and tissues (lung, lymph node, and placenta) from infected patients. Using this technology, we observe distinct subcellular localization patterns of the ORF1a and N regions, with the ORF1a concentrated around the nucleus and the N showing a diffuse distribution across the cytoplasm. In human lung tissue, we performed multiplexed RNA FISH HCR for SARS-CoV-2 and cell-type specific marker genes. We found viral RNA in cells containing the alveolar type 2 (AT2) cell marker gene (*SFTPC*) and the alveolar macrophage marker gene (*MARCO*), but did not identify viral RNA in cells containing the alveolar type 1 (AT1) cell marker gene (*AGER*). Moreover, we observed distinct subcellular localization patterns of viral RNA in AT2 cells and alveolar macrophages, consistent with phagocytosis of infected cells. In sum, we demonstrate the use of RNA FISH HCR for visualizing different RNA species from SARS-CoV-2 in cell lines and FFPE autopsy specimens. Furthermore, we multiplex this assay with probes for cellular genes to determine what cell-types are infected within the lung. We anticipate that this platform could be broadly useful for studying SARS-CoV-2 pathology in tissues as well as extended for other applications including investigating the viral life cycle, viral diagnostics, and drug screening.

## Introduction

The ongoing Coronavirus Disease 2019 (COVID-19) pandemic is caused by the betacoronavirus, Severe acute respiratory syndrome coronavirus 2 (SARS-CoV-2)^1^. COVID-19 manifests in a highly variable manner from person to person, with some infected individuals being completely asymptomatic, while others experience symptoms ranging from mild upper respiratory disease to severe pneumonia to multiorgan failure^1–3^. With such a diverse array of disease symptoms, characterizing the distribution of the SARS-CoV-2 virus across various human tissues is crucial to improving our understanding of COVID-19 pathogenesis, pathophysiology, and identifying and rationally designing effective therapies.

Critical to defining the distribution of the SARS-CoV-2 virus in humans is determining which organs and cell types become infected with SARS-CoV-2. Several studies^4–9^ have made predictions on the tissues and cell types infected by SARS-CoV-2 based on host expression of factors known to facilitate viral entry into the host cell for closely related betacoronaviruses such as SARS-CoV-1 and MERS-CoV. While this approach has helped to narrow down targets in humans, the confirmation of these predicted organs and cell types as true targets of SARS-CoV-2 remains an ongoing process. Accordingly, in a limited set of human autopsy studies, SARS-CoV-2 components (RNA and proteins) have been detected in multiple organs and organ systems including the upper airway^10,11^, lung^11,12^, gastrointestinal tract^13^, placenta^14,15^, spleen^16^, myocardium^17^, and lymph node^16^, using different combinations of RT-PCR, immunostaining, electron microscopy and *in situ* hybridization techniques. A number of these studies that use techniques with single-cell resolution (such as immunostaining and *in situ* hybridization) have identified alveolar type 2 (AT2) cells and alveolar macrophages in the lung^11^, glandular epithelial cells in the gastrointestinal tract^13^, cytotrophoblasts and syncytiotrophoblasts in placenta^18^ and macrophages in the spleen and lymph nodes^16^ as specific cell types containing SARS-CoV-2 viral components. Ultimately, defining the full range of SARS CoV-2 tropism requires direct detection approaches to validate the predictions from bioinformatic analyses in large sets of human tissue samples from COVID-19 patients.

Similarly, our current knowledge about viral life cycle and sites of SARS CoV-2 RNA synthesis is incomplete and largely based upon studies from SARS CoV-1 and MERS CoV. Upon cellular infection with these viruses, one of the early stages of the viral life cycle is assembly of the replication/transcription complex (RTC), which is the site of both viral replication and transcription of subgenomic mRNAs. Furthermore, coronaviruses both replicate their genomic RNA and transcribe subgenomic mRNAs. The subgenomic mRNAs are generated by discontinuous transcription that generates transcripts containing a conserved upstream leader sequence and a downstream body encoding viral proteins^19^. While these transcripts have been documented by Northern blot (for 40 years) and captured with sequencing techniques, they are yet to be directly visualized *in situ* in single cells.

RNA fluorescence *in situ* hybridization (FISH) techniques are ideal to address questions both about the cell types infected by SARS-CoV-2 and the subcellular localization of viral transcripts. Previously, robust RNA *in situ* assays have been developed for a number of different viral targets^20–22^. However, most human specimens from COVID patients are preserved with formalin fixation and paraffin embedding (FFPE) for long-term storage and biosafety. FFPE preserved tissues are not well-suited for single-molecule RNA FISH as the probes generate relatively low signal and the tissues have substantial autofluorescence. Furthermore, single-molecule RNA FISH probes require large regions of unique target sequence, and thus, are not amenable to specifically targeting subgenomic mRNAs, which have largely the same sequence as the genomic transcripts.

Herein, we present a platform and methodology for addressing these emerging questions about SARS-CoV-2 subcellular localization and cellular tropism using RNA fluorescence *in situ* hybridization (RNA FISH). We leverage single-molecule RNA FISH^23^ and the signal amplification capabilities of hybridization chain reaction v3.0 (HCR)^24^ to image different SARS-CoV-2 viral RNA species in cell culture infection models and FFPE human autopsy specimens. We extend the assay to multiplex probe sets for viral and host RNAs to simultaneously detect cells with viral RNA and determine their cell type. This platform allows us to observe differences in RNA staining patterns of SARS-CoV-2 infection between AT2 cells and alveolar macrophages in human lung autopsy tissue.

## Results

RNA *in situ* hybridization technologies offer the ability to visualize RNA within fixed cells and tissues. Such technologies have been used for both cellular RNAs and viral RNAs in infected cells^21-23,25,26^. With the emergence of the SARS-CoV-2 virus, we sought to use RNA *in situ* hybridization techniques to visualize the viral RNA transcripts in both cell lines and tissues. To overcome limitations from background autofluorescence and for robust RNA detection, we used hybridization chain reaction v3.0 (HCR) to achieve amplification of the RNA FISH signal^24^. We developed probe sets consisting of multiple probe pairs that are tiled along the RNA sequence of interest. Each probe pair, termed “split-initiator” probes, contains a region of complementarity to the viral RNA and half of the initiator sequence for signal amplification via polymerization of dye-conjugated DNA hairpins. Because the initiator sequence is divided between the two split-initiator probes, amplification only occurs if both of the probes bind adjacently, providing additional specificity for the target of interest (Fig. 1a). As previously described^24^, we used a two-stage protocol in which we first hybridize the split-initiator probes and then amplify the signal using fluorescently labeled DNA hairpins. We can multiplex the assay for multiple targets by using distinct hairpin sequences labeled with different fluorophores for each RNA target.

**Figure 1:**
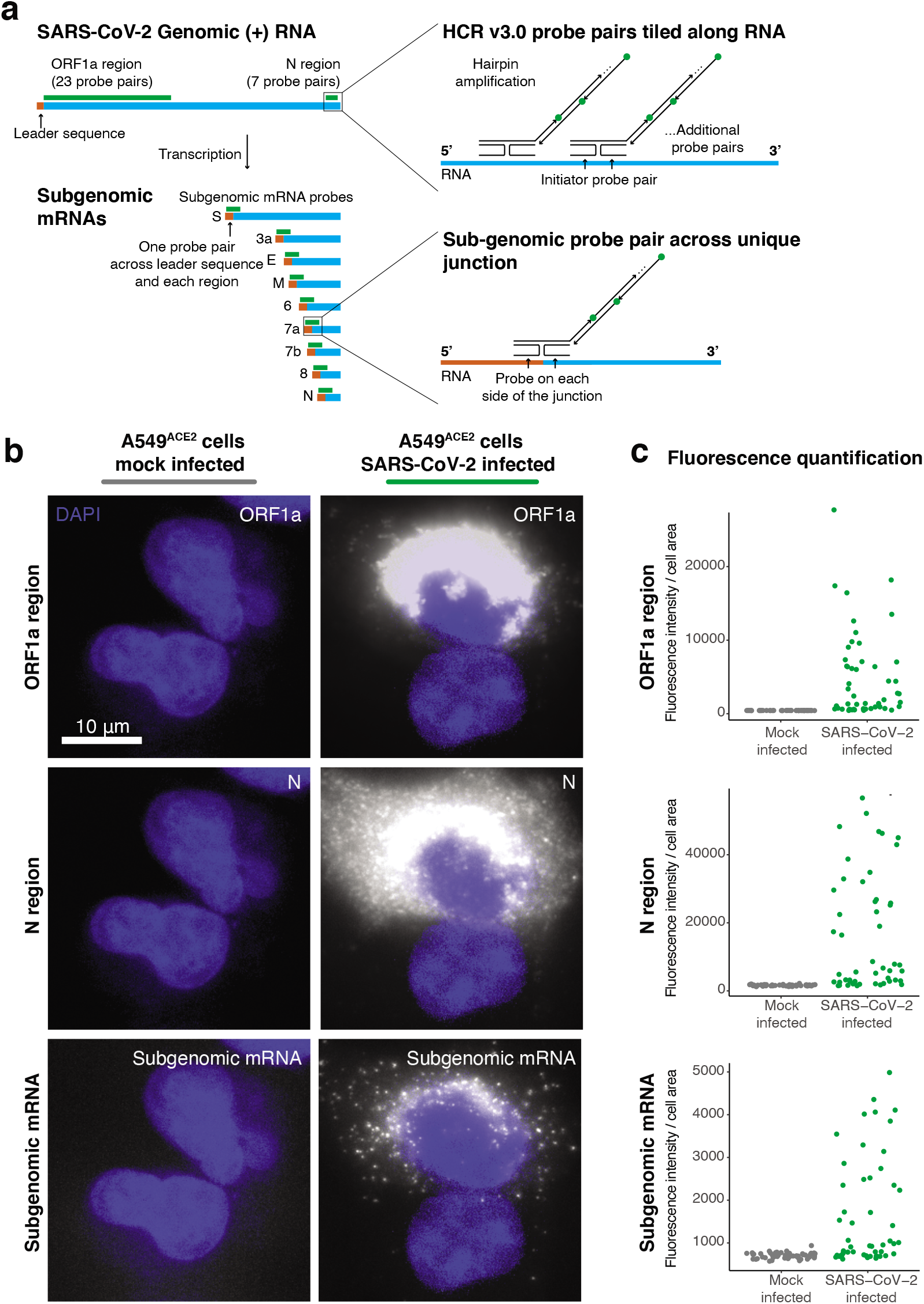
RNA FISH HCR v3.0 probe sets enable direct visualization of the SARS-CoV-2 virus. **a.** Schematic of the SARS-CoV-2 genomic RNA and subgenomic RNA species with HCR v3.0 probe designs highlighted. We developed probes tiled along the ORF1a and N regions of the SARS-CoV-2 (+) RNA strand. These probe sets consisted of 23 probe pairs for ORF1a and 7 probe pairs for N. To detect all of the sub-genomic RNAs, we positioned the HCR probes across the junction of the leader sequence and each unique sub-genomic transcript. In the schematic, the leader sequence is shown in orange, the transcript is shown in blue and the probe design is shown in green. **b.** Representative images of the A549^ACE2^ cells mock infected or infected with SARS-CoV-2 at an MOI=1, fixed 24 hours post infection, and then hybridized with probes for ORF1a, N, and sub-genome. DAPI labels cell nuclei. Scale bar applies to all images and shows 10 μm. **c.** Quantification of the fluorescence signal from the experiment in b. For each, the mock infected data set (shown in gray) and the SARS-CoV-2 infected data set (shown in green), we quantified fluorescence signal intensity from 50 cells per condition. Note that the SARS-CoV-2 infected sample contained both cells that were infected and cells that remained uninfected.

We first designed probe sets targeting the positive stranded SARS-CoV-2 RNA sequences of the ORF1a and N regions (Fig. 1a). We tested these probe sets in A549^ACE2^ human lung cancer cells infected with SARS-CoV-2 at a multiplicity of infection (MOI) of 1, as well as mock-infected A549^ACE2^ cells. We found high intensity fluorescent labelling with both the ORF1a and N probe sets in the infected, but not in mock-infected samples (Fig. 1b). Staining from both probe sets was confined to the cytoplasm of cells and did not stain inside the cell nucleus. Interestingly, we observed that the ORF1a probe set (which labels only genomic RNA) showed the highest intensity of staining in a region around the periphery of the nucleus of each cell. This staining pattern is consistent with reported coronavirus RNA replication at replication/transcription complexes (RTCs), which are networks consisting of host endoplasmic reticulum-derived, perinuclear, double-membrane structures^27–29^. Meanwhile, the N region probe set showed more diffuse staining throughout the cytoplasm, but higher intensity in the perinuclear region. Such a pattern could be expected for the N region probes as they are likely binding both genomic RNA species in the RTCs and all the subgenomic mRNA species (Fig. 1a). These subgenomic mRNAs are translated by the host ribosome, and thus are more diffuse through the cytoplasm rather than largely confined to viral replication centers.

To further resolve the localization of genomic and subgenomic mRNA, we designed probes to uniquely label the subgenomic mRNA species without simultaneously targeting SARS-CoV-2 genomic RNA. Such a probe design is difficult because the subgenomic mRNA sequence is also contained within the genomic RNA, and thus subgenomic and genomic transcripts would be simultaneously targeted with conventional probe designs. Thus, to develop subgenome specific probes, we leveraged a feature of coronavirus transcription biology. To generate subgenomic mRNAs, the viral polymerase first transcribes negative-strand intermediates from which it then transcribes the subgenomic mRNAs. During this synthesis of negative-strand intermediates, the polymerase terminates transcription when it encounters transcription regulatory sequences (TRSs) upstream of each subgenomic mRNA open reading frame and resumes at a TRS located further towards the 5’ end of the genomic template. This interrupted form of transcription, known as discontinuous transcription^19,30^, adds an antisense copy of the genomic leader sequence to each subgenomic mRNA intermediate. Therefore, in the subgenomic mRNAs only, there is a unique junction formed between the 3’ end of the leader sequence and the 5’ end of their gene sequence. To target each individual subgenomic mRNA, we designed HCR probe pairs that span the unique junction sites, with one of the split-initiator probes positioned on the leader sequence and the other split-initiator probe on the gene sequence (Fig. 1a). Because each split-initiator probe contains half the initiator sequence, amplification should only occur if the two probes bind adjacent to each other, which would be the case for each target subgenomic mRNA but not genomic RNA. With this strategy, we achieve highly specific detection of the fusion transcripts containing the leader and each subgenomic sequence. We designed these subgenomic probes for each of the eight different canonical subgenomic mRNA species and used them together on A549^ACE2^ cells infected with SARS-CoV-2 (Fig. 1b). We found that the subgenomic mRNA probe sets showed a diffuse staining pattern throughout the cytoplasm that is distinct from both the ORF1a and N probe sets. This diffuse staining pattern is consistent with the subgenomic mRNAs being distributed throughout the cytoplasm for translation by free ribosomes or ER associated ribosomes (rather than concentrated in the replication/transcription complex as seen for the ORF1a probe set).

Next, we quantified the fluorescence intensity from each of these probe sets in both the infected and mock-infected cells. We found that the mean fluorescence intensity of infected cells was significantly higher than the fluorescence intensity of uninfected cells in the mock infected sample (Fig. 1c). Of note, the infected samples contained a mixture of infected and non-infected cells, which are clearly distinguishable *in-situ*. We also designed conventional (non-amplified) single-molecule RNA FISH probes to the same regions of SARS-CoV-2 genomic RNA and compared the signal from the amplified RNA FISH HCR to the non-amplified single-molecule RNA FISH (Supp. Fig. 1). We again found with single-molecule RNA FISH that the ORF1a probe set had a perinuclear staining pattern, while the N probe had a more diffuse cytoplasmic distribution. Compared to non-amplified single-molecule RNA FISH, the RNA FISH HCR signal was significantly brighter, requiring much shorter exposure times for imaging. Thus, we selected RNA FISH HCR for subsequent experiments in tissues in which we expected higher background compared to cell culture conditions.

We next sought to use this assay in human tissues to localize sites of SARS-CoV-2 viral RNA. We analyzed human autopsy specimens from research autopsies of COVID-19 patients performed at the University of Pennsylvania and Children’s Hospital of Philadelphia in 2020. The tissues were fixed in neutral buffered formalin, embedded in paraffin, sectioned, and then probed for SARS-CoV-2 RNA (Fig. 2a). We analyzed a total of 14 different lung specimens from eight patients and found one case that showed extensive staining of SARS-CoV-2 RNA in lung tissue (Supp. Table 1). Of note, the patient with extensive virus staining in the lung was immunosuppressed, and decompensated within 2 days of arriving at the hospital. In this specimen, we observed discrete regions of the lung containing infected cells (Fig. 2b, Lung) as well as regions with bright fluorescence signal, but no nuclei present (Supp. Fig 2a). To find areas with SARS-CoV-2 staining, we developed a computational pipeline (described in methods) to segment cells and then quantify the fluorescence staining from the ORF1a viral probe set (output of the analysis is shown in Supp. Fig. 3). As a control, we also examined lung tissue from patients who were not infected with SARS-CoV-2. In these controls, our computational analysis did not identify any cells passing the threshold of significant ORF1a viral RNA staining (Supp. Fig. 4).

**Figure 2:**
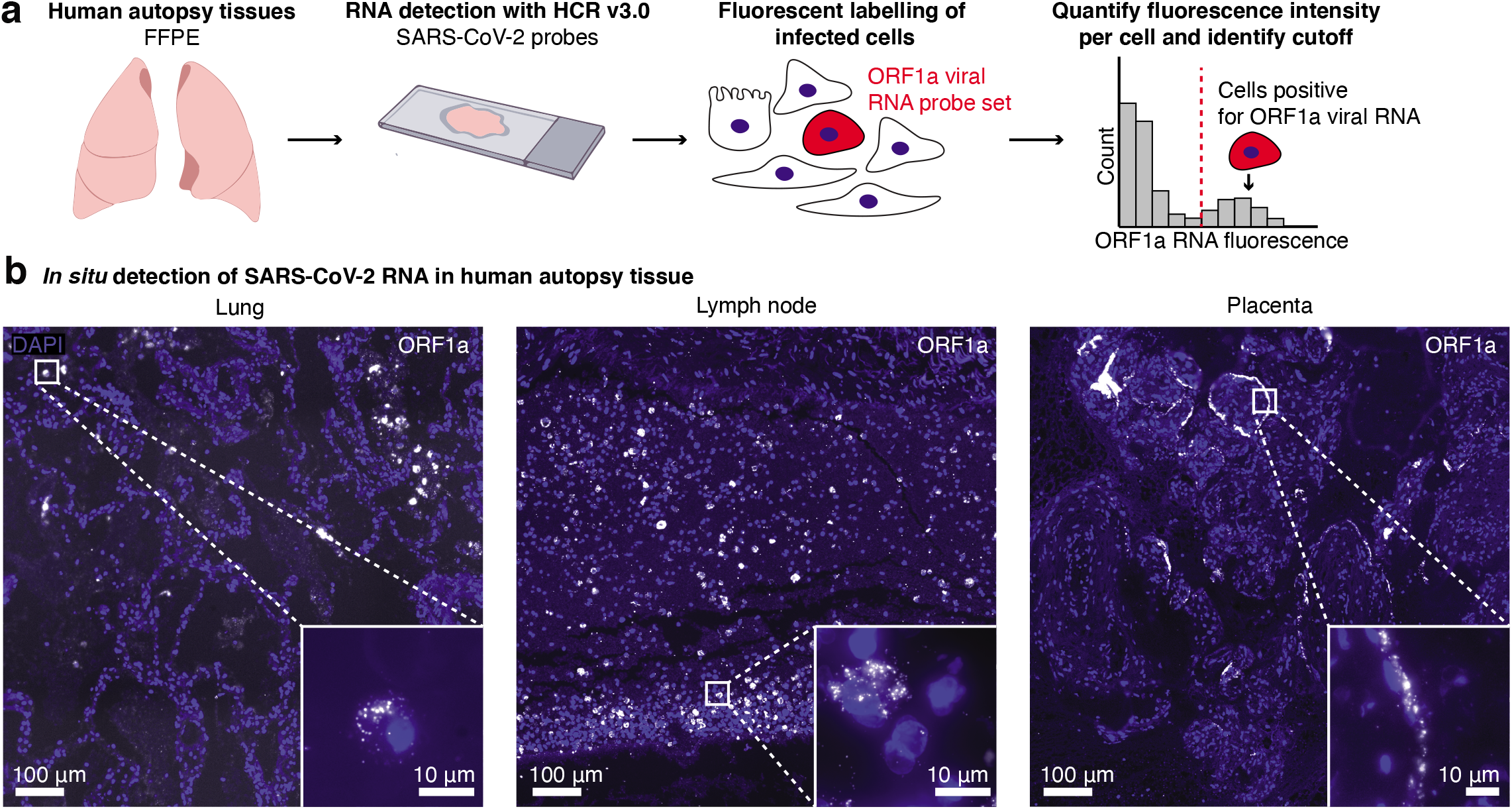
RNA FISH HCR in FFPE human autopsy tissues. **a.** Experiment design in which we performed RNA FISH HCR with ORF1a probe sets on FFPE tissues including lung, hilar lymph node, and placenta. **b.** Example images of each tissue with ORF1a RNA staining. Images are large area scans of image tiles acquired at 20X. Scale bar on the large images shows 100 μm. Inset images show a zoomed in example of ORF1a RNA staining in that tissue. Scale bars on these inset images are 10 μm. DAPI stain (blue) labels the cell nuclei in all images.

From the same patient with extensive lung infection, we surveyed other tissues for SARS-CoV-2 RNA. We performed RNA FISH HCR with the ORF1a virus probes on a total of eleven different tissues including esophagus, kidney, liver, hilar lymph node, spleen, heart, stomach, ileum, duodenum, jejunum and trachea. Of all of these tissues, we only detected viral RNA with our probe sets in the hilar lymph node. Of note, it is possible that some of the other tissues from this patient also contained virus, but underwent more degradation prior to RNA FISH HCR. In the lymph node specimen, we found ORF1a RNA FISH HCR signal localized to cells scattered throughout the tissue (Fig. 2b). These results are consistent with other studies reporting the detection of SARS-CoV-2 nucleocapsid-positive cells in hilar lymph nodes^31,32^.

We also analyzed two human placenta samples from cases in which the mother tested positive for SARS-CoV-2. Both samples showed cells with ORF1a probe set staining localized predominantly along the periphery of villi structures (Fig. 2b, Placenta). This pattern is consistent with other reports using immunohistochemical assays, electron microscopy, and RNAscope *in situ* hybridization^18,33-35^ that show viral localization to syncytiotrophoblasts, which are located along the villous periphery and interface with maternal blood. We further confirmed this observation through comparison with an adjacent hematoxylin and eosin (H&E) stained slide (Supp. Fig. 5).

After using these probe sets to localize the virus across tissues, we next wanted to know whether multiplexed RNA FISH HCR could be used to determine what cell types become infected with the virus in the lung. We developed a robust strategy for selecting cell type specific marker genes for multiplexed *in situ* analysis along with SARS-CoV-2 RNA. We used single-cell RNA-sequencing data from the human lung atlas^36^ to identify genes that would uniquely label alveolar type 1 (AT1) cells, alveolar type 2 (AT2) cells, and alveolar macrophages within the lung. For our analysis, we considered the specificity of the marker gene, the fraction of cells that expressed the marker, and the expression level (as we needed genes with high enough expression for accurate detection *in situ*). We only considered markers that were present in all of the human lung cell atlas subjects to avoid genes with heterogeneous expression between individuals. We developed HCR probes for 1 marker of each cell type (*AGER* for AT1 cells, *SFTPC* for AT2 cells, and *MARCO* for alveolar macrophages, Supp. Fig. 6).

We first performed multiplexed RNA FISH HCR on lung autopsy tissue using probe sets for *AGER* (to mark AT1 cells), *SFTPC* (to mark AT2 cells) and SARS-CoV-2 ORF1a RNA (Fig. 3a). We observed *AGER-* and *SFTPC*-high cells throughout the tissue (Fig. 3c). Reassuringly, we did not find many cells showing high levels of both genes (only 1 observed), allowing us to identify *AGER-* and *SFTPC*-high cells as AT1 and AT2 cells, respectively. Furthermore, the cells labeled by the *AGER* and *SFTPC* probe sets had morphologies consistent with AT1 and AT2 cells, respectively. For each marker gene, we selected a threshold number of mRNA molecules per cell to identify cells as either AT1 or AT2 cells (described in methods). We then analyzed the fluorescence intensity of the ORF1a SARS-CoV-2 probe set signal across all 252,820 cells in the dataset from patient 2 with high levels of infection in the lung. We found that a large fraction of the *SFTPC*-positive AT2 cells also stained as positive for ORF1a RNA (27.4%) but that none (0%) of the *AGER*-positive AT1 cells were positive for SARS-CoV-2 (Fig. 3b,d). There were also a substantial number of cells (647) that were ORF1a RNA positive, but did not have expression of either *SFTPC* or *AGER*. Additionally, there were a number of cells with viral RNA staining that lacked nuclear DAPI staining suggesting that these represent dead cells that were previously infected (Supp. Fig. 2b). In sum, we demonstrated highly specific discrimination of AT1 and AT2 cells, and found that only the AT2 cell population had robust SARS-CoV-2 RNA consistent with these cells being the major target of infection

**Figure 3:**
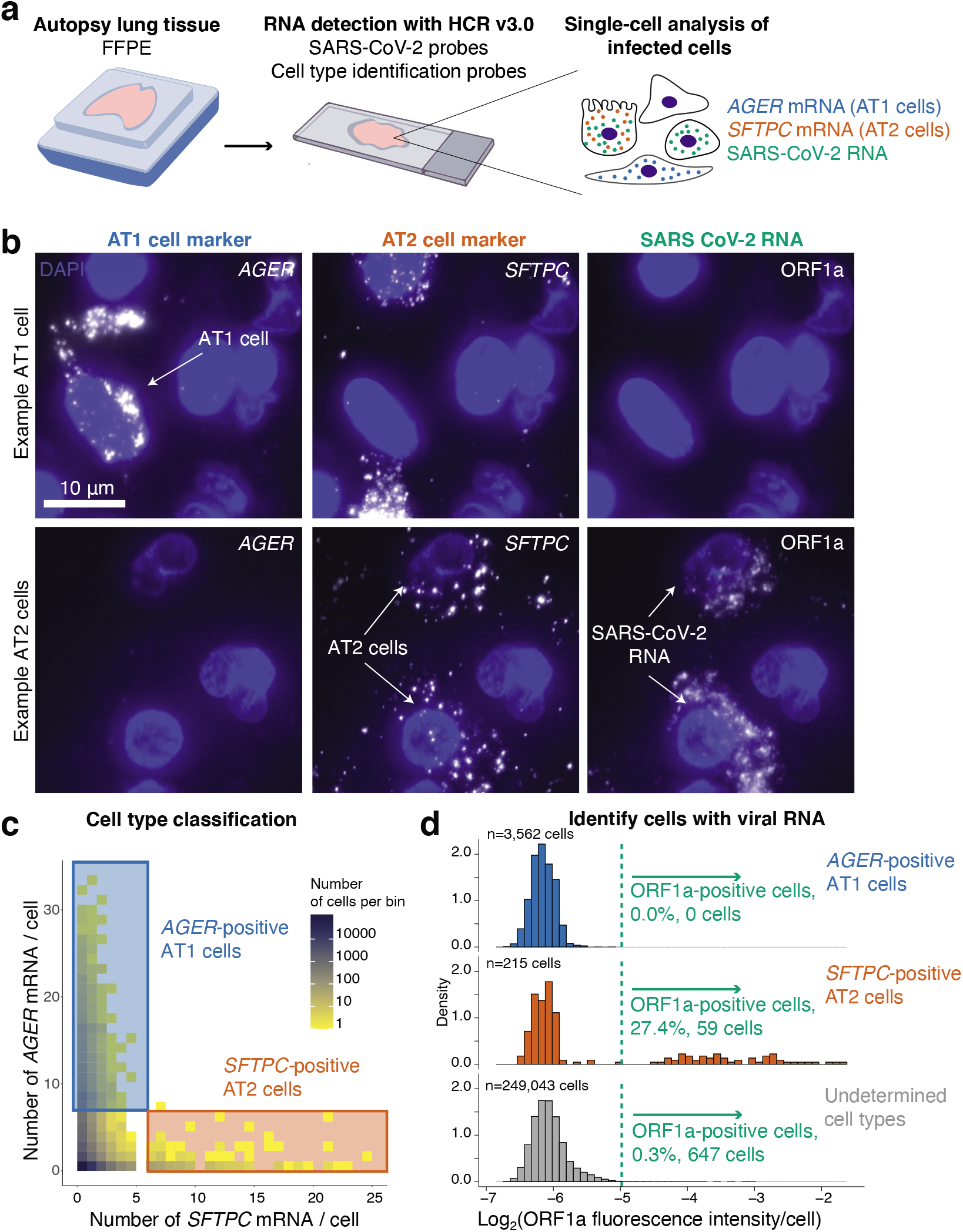
Multiplexed RNA FISH HCR identifies AT2 cells containing viral RNA in lung autopsy samples. **a.** We probed FFPE human lung tissue with SARS-CoV-2 probe sets as well as probe sets for cell-type specific marker genes, *AGER* for AT1 cells and *SFTPC* for AT2 cells. **b.** Representative images of cells classified as AT1 cells or AT2 cells. Top row depicts an AT1 cell staining positive for *AGER*. Second row shows two *SFTPC*-positive AT2 cells staining with the ORF1a viral RNA probe set. DAPI stain (blue) labels the cell nuclei in all images. Scale bars show 10 μm. **c.** Here, we acquired large tiled image scans consisting of 252,820 cells total. We quantified the *AGER* and *SFTPC* mRNA in each cell and set a cutoff (see methods) for determining which cells are positive for each gene indicating that they are either AT1 or AT2 cells, respectively. Plot shows a scatterplot of mRNA levels with cutoffs for AT1 and AT2 cells. The color on the scatterplot indicates the number of cells at each point on the plot and the scale is shown by the legend with yellow being low cell numbers and blue high cell numbers. The blue rectangle shows the region on the plot for *AGER*-positive AT1 cells and the red rectangle shows the region on the plot for *SFTPC*-positive AT2 cells. **d.** Histograms of the log2 of the fluorescence intensity for the SARS-CoV-2 ORF1a probe set in each cell. The data is split into three histograms for each cell identified (AT1 cells, AT2 cells, undetermined cells). The green dotted line shows the cutoff for calling a cell as postivie for ORF1a viral RNA.

We next sought to determine what other cell types contain SARS-CoV-2 RNA in the lung. We multiplexed a probe set for a macrophage specific gene (*MARCO*) with the SARS-CoV-2 ORF1a probe set (Fig. 4a). We found a large number of *MARCO*-positive cells that contained staining with the SARS-CoV-2 ORF1a probe set (Fig 4b). However, in many of these cells, the subcellular localization of the signal was distinct from the staining that we previously observed in *SFTPC*-positive AT2 cells (Fig. 4c). In many examples, the staining within alveolar macrophages appeared to be compartmentalized within a smaller region of the cytoplasm (compared to the AT2 cells). In AT2 cells, the ORF1a virus probe set typically stained the entire cell cytoplasm and around the entire periphery of the nucleus. It is possible that in the alveolar macrophages these smaller regions of staining represent restriction of the viral RNA to a subcellular compartment such as the phagolysosome. This difference in RNA localization suggests that viral RNA enters alveolar macrophages through phagocytosis of other infected cells or infected debris, which has been suggested by others^37–39^.

**Figure 4:**
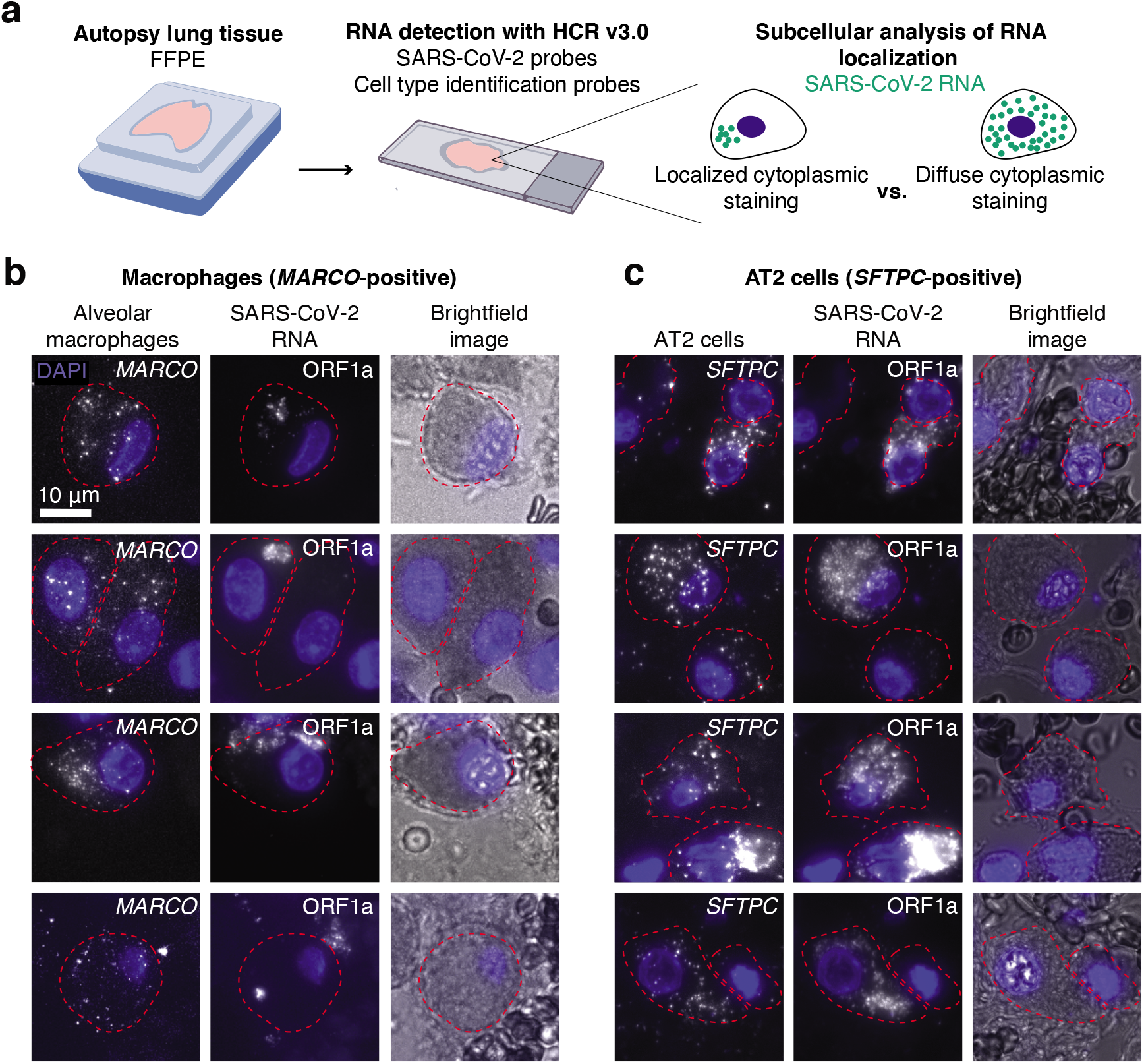
Alveolar macrophages and AT2 cells show distinct viral RNA staining patterns in autopsy tissue. **a.** Schematic of experimental design in which we multiplexed cell-type specific marker genes with SARS-CoV-2 ORF1a probes. We examined the subcellular distribution of viral RNA staining in alveolar macrophages and AT2 cells. **b.** Examples of alveolar macrophages showing *MARCO*, ORF1a, and brightfield images for each cell. The border of each cell’s cytoplasm is shown by the red dotted line in each image. DAPI stain for cell nuclei shown in blue. Scale bars show 10 μm. **c.** Examples of AT2 cells showing *SFTPC*, ORF1a, and brightfield images for each cell. The borders, nuclei, and scale bars are labeled the same as in B.

## Discussion

In this paper, we outline methods and probe designs for visualizing SARS-CoV-2 RNA in cell lines and human autopsy specimens. First, to enable compatibility with autopsy tissue from COVID-19 patients, we developed a protocol for tissue processing (described in methods). Next, to tailor the probes to the SARS-CoV-2 virus, we designed unique probe sets for the ORF1a and N region RNA. We further developed the assay to be able to uniquely label subgenomic mRNAs, which is not possible with conventional probe designs. We validated each of these probe sets in cell culture models and then applied them to autopsy tissues. We identified infected cells as those labeled by ORF1a RNA in lung, lymph node, and placenta. Finally, we performed multiplex RNA FISH HCR with probe sets for cell-type specific marker genes to determine that AT2 cells and alveolar macrophages contain SARS-CoV-2 RNA in the lung. Through subcellular visualization of the RNA localization, we found that the subcellular localization of ORF1a-containing transcripts is different between AT2 cells and alveolar macrophages. This finding further supports previous studies suggesting that alveolar macrophages acquire SARS-CoV-2 RNA through phagocytosis rather than receptor mediated entry^37,38^. Taken together, these technological advances in RNA FISH HCR present a powerful toolset for visualizing SARS-CoV-2 in cell culture and FFPE tissues.

The primary alternative methods for staining viruses in tissues rely on antibodies including immunohistochemistry and immunofluorescence. In the setting of a pandemic with a new virus, the speed of antibody development, which can take weeks to months, can present significant challenges. Furthermore, antibody development can be costly, and even after production, antibodies still require extensive validation to prove that they are correctly targeting the protein of interest. In contrast, with modern sequencing-based epidemiologic surveillance, a novel agent’s genome may be available in days, and simple rules govern design of suitable hybridization probes. Thus, it is cheaper, easier, and faster to develop RNA probes using oligonucleotides. Since IHC/IF and RNA FISH target different molecules (protein vs. RNA, respectively), they also provide different and complementary information about the virus. With SARS-CoV-2, a number of studies have found viral protein staining by immunofluorescence or immunohistochemistry in different tissues, but have not provided sufficient evidence to confirm that there is replicating virus present^11–13^. As RNAs are more labile than proteins, one potential strength of RNA-based techniques is that they may be a more reliable readout of ongoing viral infection.

Another relevant technology for viral detection is RT-PCR, which is commonly used for SARS-CoV-2 diagnostics. The primary strength of RT-PCR is sensitivity as it can detect down to the scale of tens to hundreds of transcripts^41,42^. While sensitivity makes RT-PCR very useful for diagnostics, it requires that the entire sample is combined into one tube. With RT-PCR, different cell types are mixed, subcellular localization of RNA is lost, and the global tissue architecture is also lost. Furthermore, these techniques work better on viable or fresh frozen samples, which are often limited, as most tissue banking resources largely contain FFPE specimens. Thereby, while RNA FISH HCR is unlikely to have the same sensitivity as RT-PCR, it offers direct visualization of RNA which allows for single-cell analysis to identify infected cell types, subcellular localization of different RNA transcripts, and spatial analysis of viral and cellular RNA across infected tissues.

RNA FISH techniques have specific advantages over other tissue-based techniques for visualizing viruses. In the setting of a new viral threat, custom probes can be quickly designed to target the virus, only requiring knowledge of the viral sequence. Our probe design approach here ensures that the probes are specific only to the desired virus by querying sequence databases of other viruses. After designing the probe sequences, the synthesis of oligonucleotide probes is both inexpensive and fast such that the entire assay can easily be set up for a new virus in a relatively short period of time (3-5 days). As we do in this paper, RNA FISH-based probes can be designed to different RNA species generated by the virus and even used to discriminate between closely related virus strains through both bioinformatic probe design strategies and different probe conformations. Examples include this study in which we target both genomic RNAs and subgenomic mRNAs, and other studies with probes to positive and negative RNA strands^40^, different segments of the influenza genome^21,22^, and even probes that detect single-base pair variants within the virus^22^.

In addition to probes that target different components of the virus, we can also multiplex viral probes with probes for cellular genes. Here we target cellular genes to identify cell types within the infected tissue, but this approach could be applied to profile other genes involved in the host response to viral infection. Furthermore, with recent advances in RNA *in situ* technologies it is now possible to probe hundreds to thousands of genes with techniques such as seqFISH+ and MERFISH^41,42^. These platforms could be easily adapted to include virus probes as well. Such an approach could reveal the full picture of how a viral infection alters a tissue, including the direct effects on the cells that are infected by the virus, as well as the effects on the neighboring cells and immune response.

In summary, this paper outlines protocols and probe sets for using RNA FISH HCR in FFPE tissues for visualizing SARS-CoV-2 RNA. We demonstrate the use of these methods for visualizing different viral RNA species, identifying infected cells in FFPE tissues, and determining the cell types that are infected. Our work establishes RNA FISH HCR as a powerful technique for virology and pathology to visualize SARS-CoV-2 RNA in tissues that can be easily extended for new infectious diseases in the future.

---

**Supplementary Figure 1:**
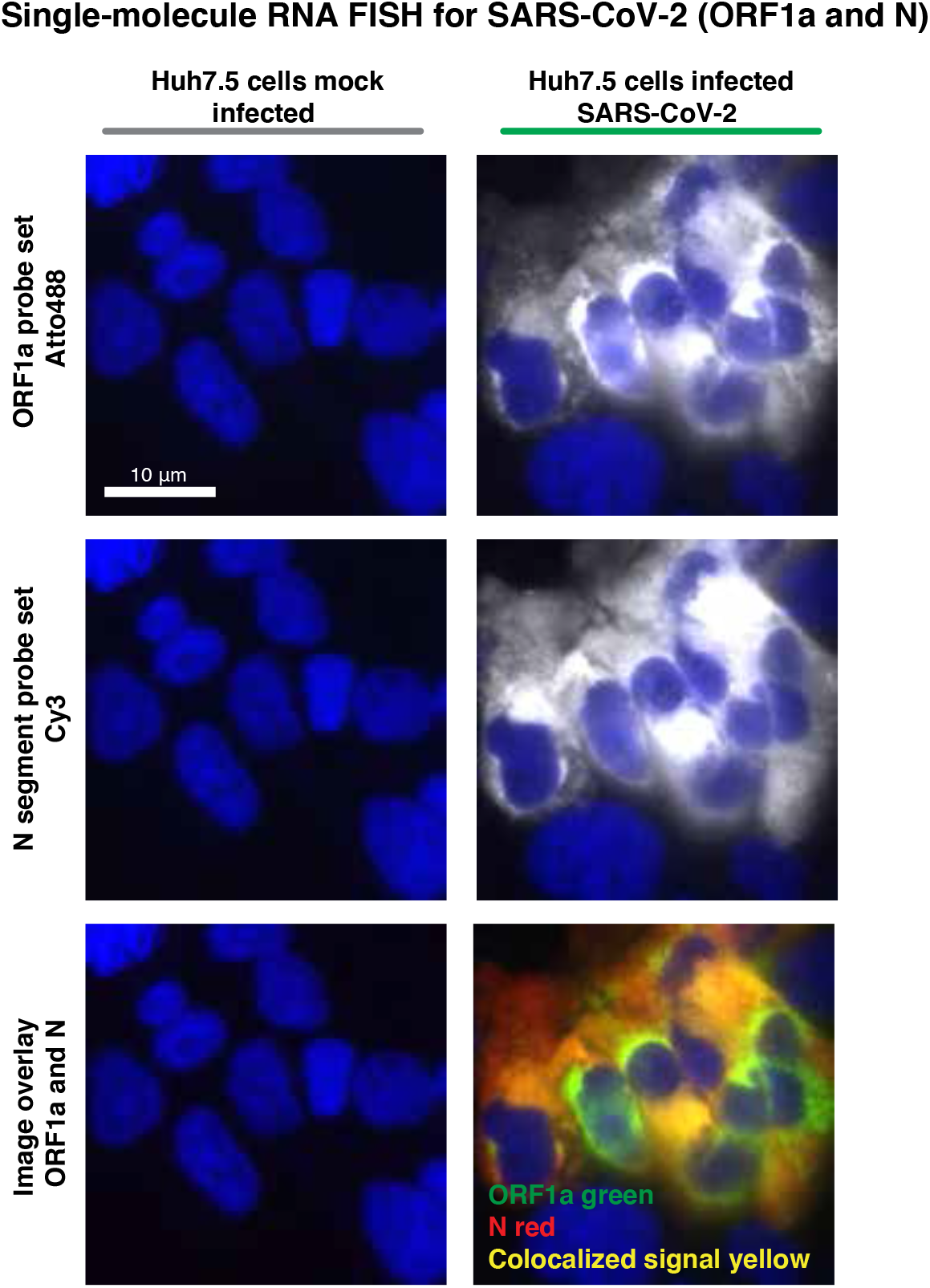
Single-molecule RNA FISH with probes targeting ORF1a and N regions in Huh7.5 cells infected with SARS-CoV-2. Representative images of Huh7.5 cells infected with SARS-CoV-2 and hybridized with RNA FISH probes targeting ORF1a and N. Similar to the RNA FISH HCR, we observed higher fluorescence signal intensity in the perinuclear region of the ORF1a probe compared to the N probe. The bottom row of images is a composite of the ORF1a probe (green), N probe (red), and DAPI signal. In all images, the DAPI stain for cell nuclei is shown in blue. Scale bars are 10 μm.

**Supplementary Figure 2:**
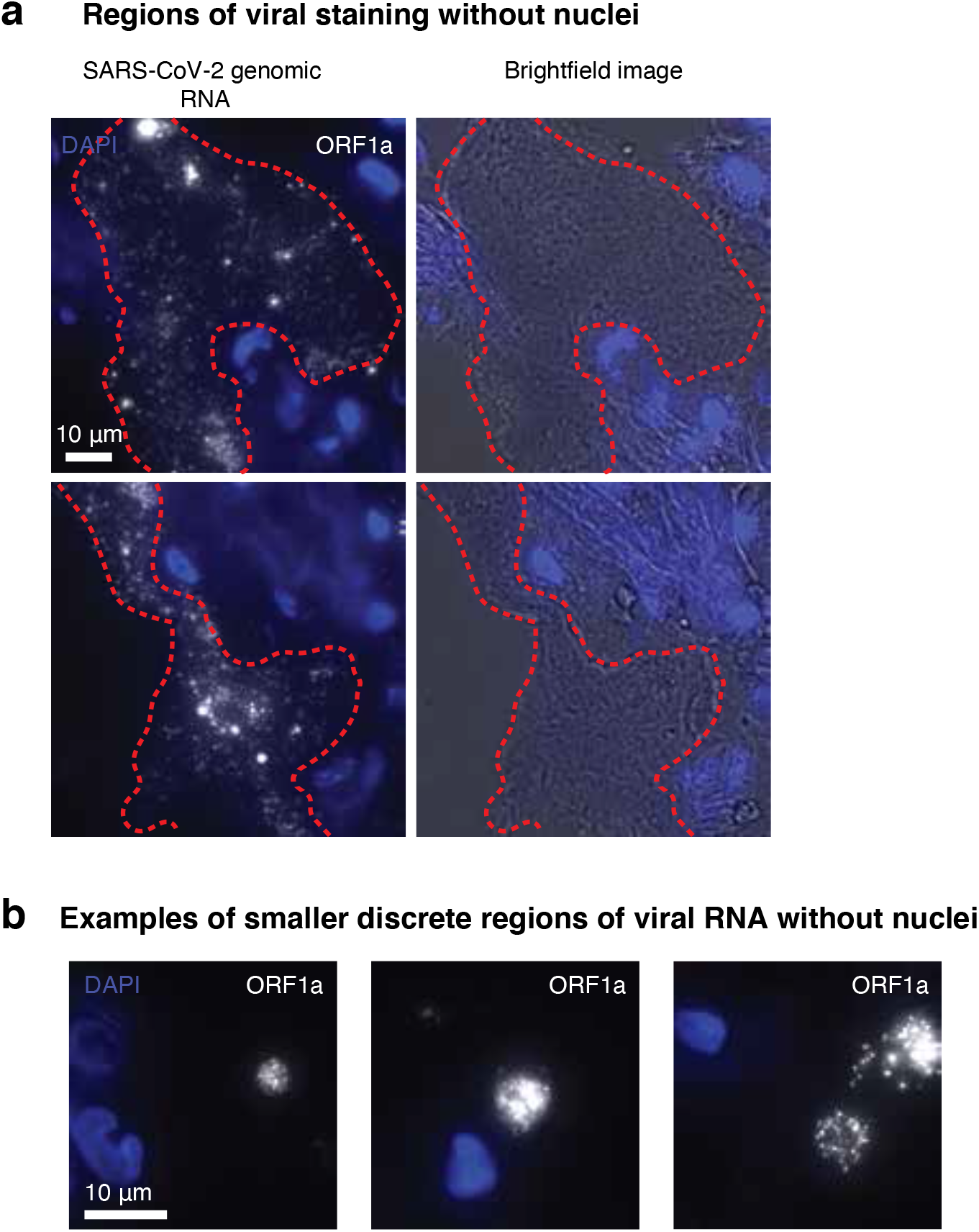
Examples of regions in the lung tissue that show viral staining with ORF1a probes, but do not have nuclei. A. Two example regions in which we observed extensive viral staining with the ORF1a probe set, but did not observe DAPI signal. Right image is RNA FISH HCR for ORF1a and the left is the brightfield image. Red dotted lines show areas of interest with ORF1a staining. B. Examples of small discrete regions of ORF1a staining without DAPI staining. In all images, the DAPI stain for cell nuclei is shown in blue. Scale bars are 10 μm.

**Supplementary Figure 3:**
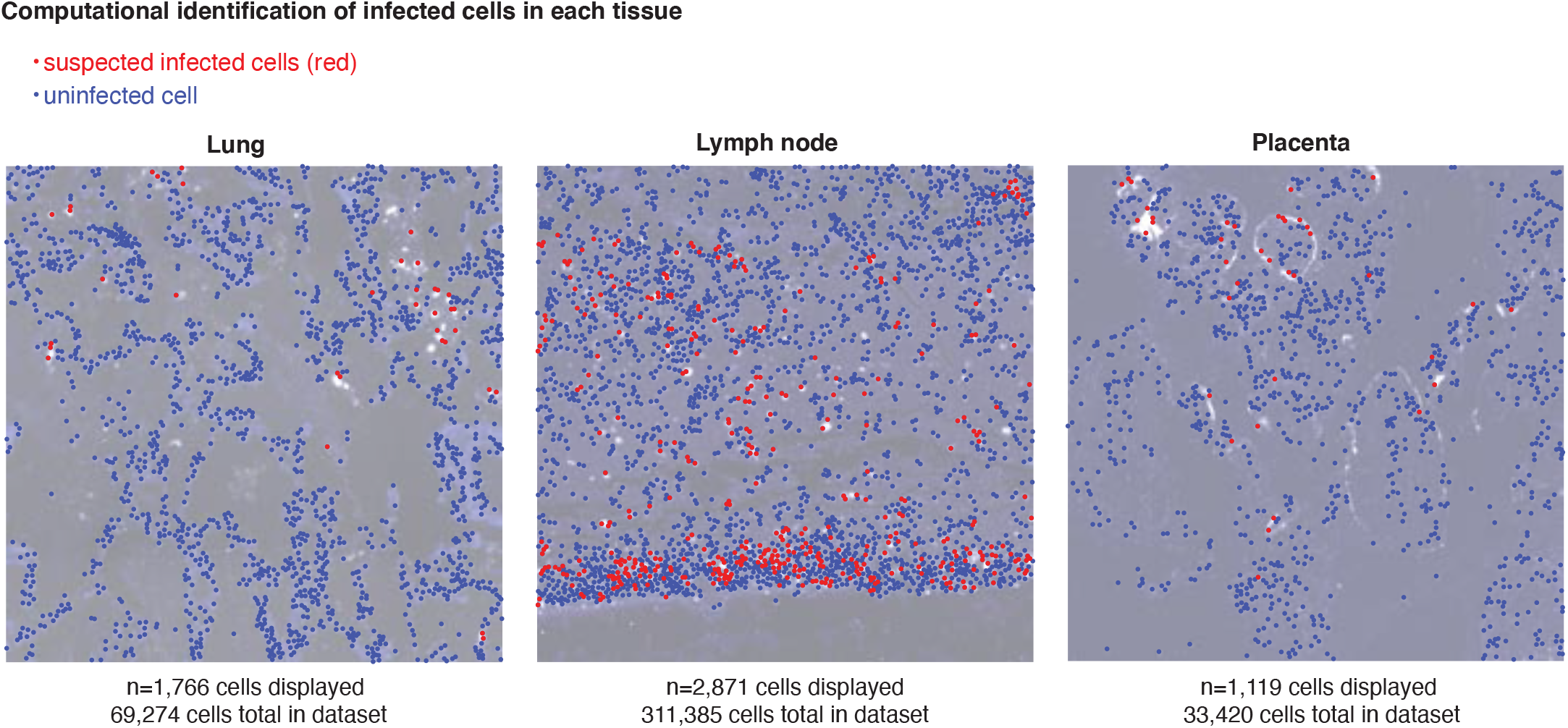
Computational analysis identifying suspected infected cells in each tissue. Images overlayed with the results from our image processing pipeline. Blue dots label cells with fluorescence signal below the cutoff for infected and red dots label cells above the cutoff for infected. Images are from a subset of the data. Total number of cells displayed in each region and the total number of cells in each data set is below each image.

**Supplementary Figure 4:**
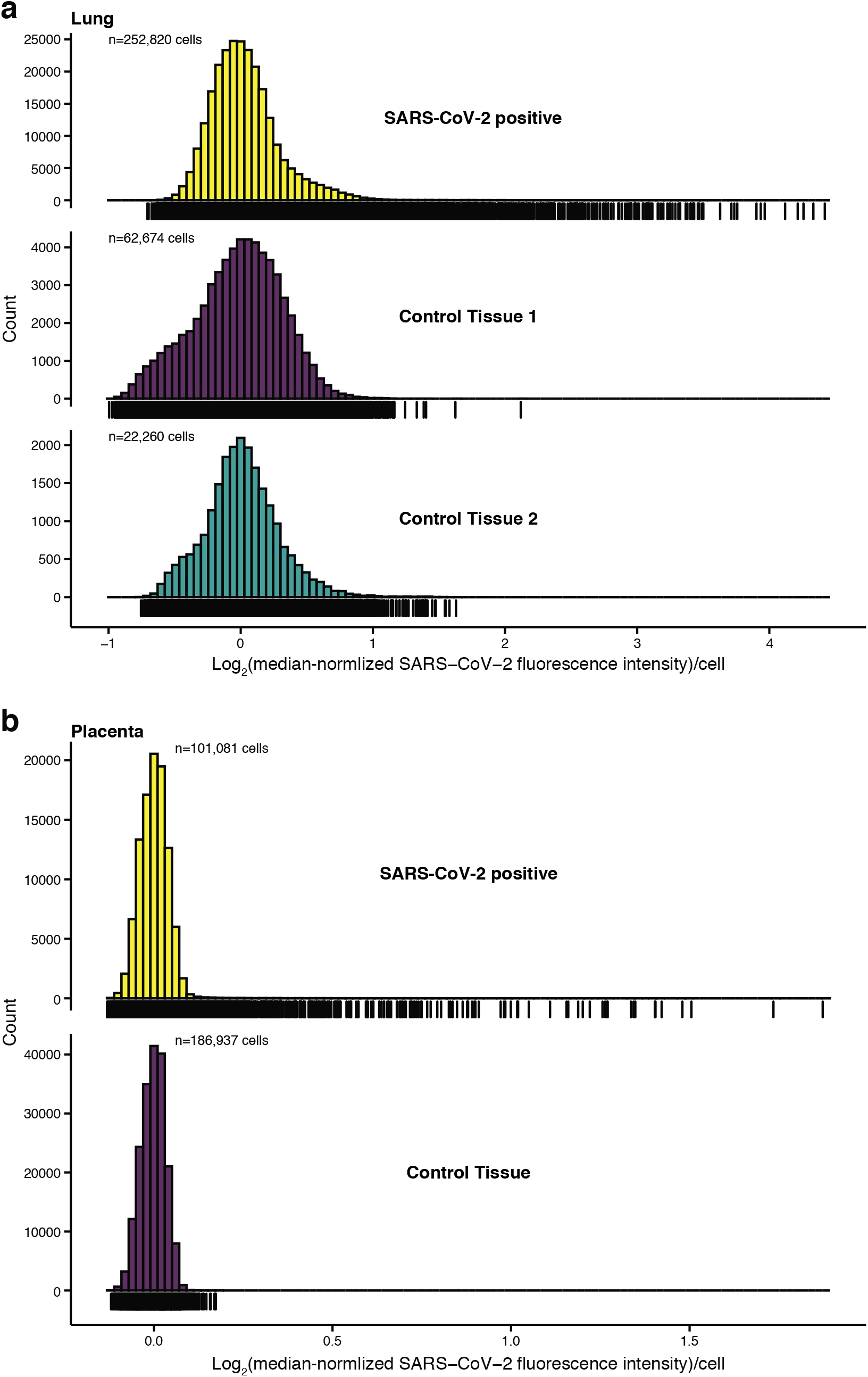
Comparison of infected tissue to control tissues samples in which the patients did not have SARS-CoV-2 infection. We performed RNA FISH HCR in A. lung and B. placenta samples from patients with SARS-CoV-2 infection (yellow) and control samples from patients that did not have SARS-CoV-2 infection (purple and teal). Histograms display the Log_2_ transformation of median normalized ORF1a fluorescence signal in each cell. Infected samples have a much higher median-normalized ORF1a fluorescence signal than control tissue.

**Supplementary Figure 5:**
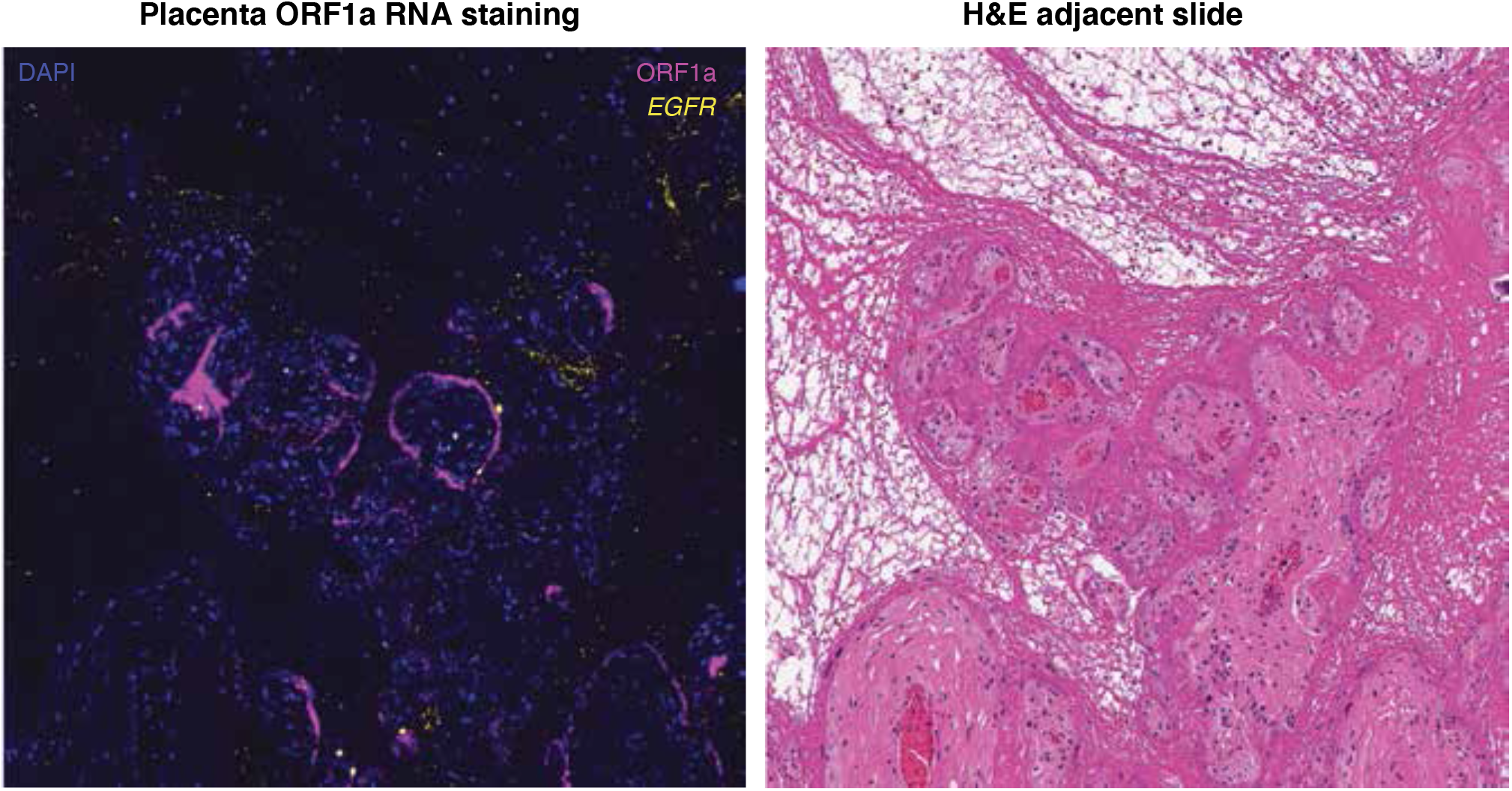
Example region of placenta with RNA FISH HCR for ORF1a with an adjacent tissue section stained with H&E. We performed RNA FISH HCR with probe sets for ORF1a and EGFR. On the adjacent section, we stained the tissue with hematoxylin and eosin. We took tiled image scans of the fluorescence slide and used a slide scanner for the H&E. We aligned the two images to identify the corresponding H&E region for which we found cells staining with the ORF1a probe set. ORF1a fluorescence signal is in pink, *EGFR* is in yellow, and DAPI is in blue.

**Supplementary Figure 6:**
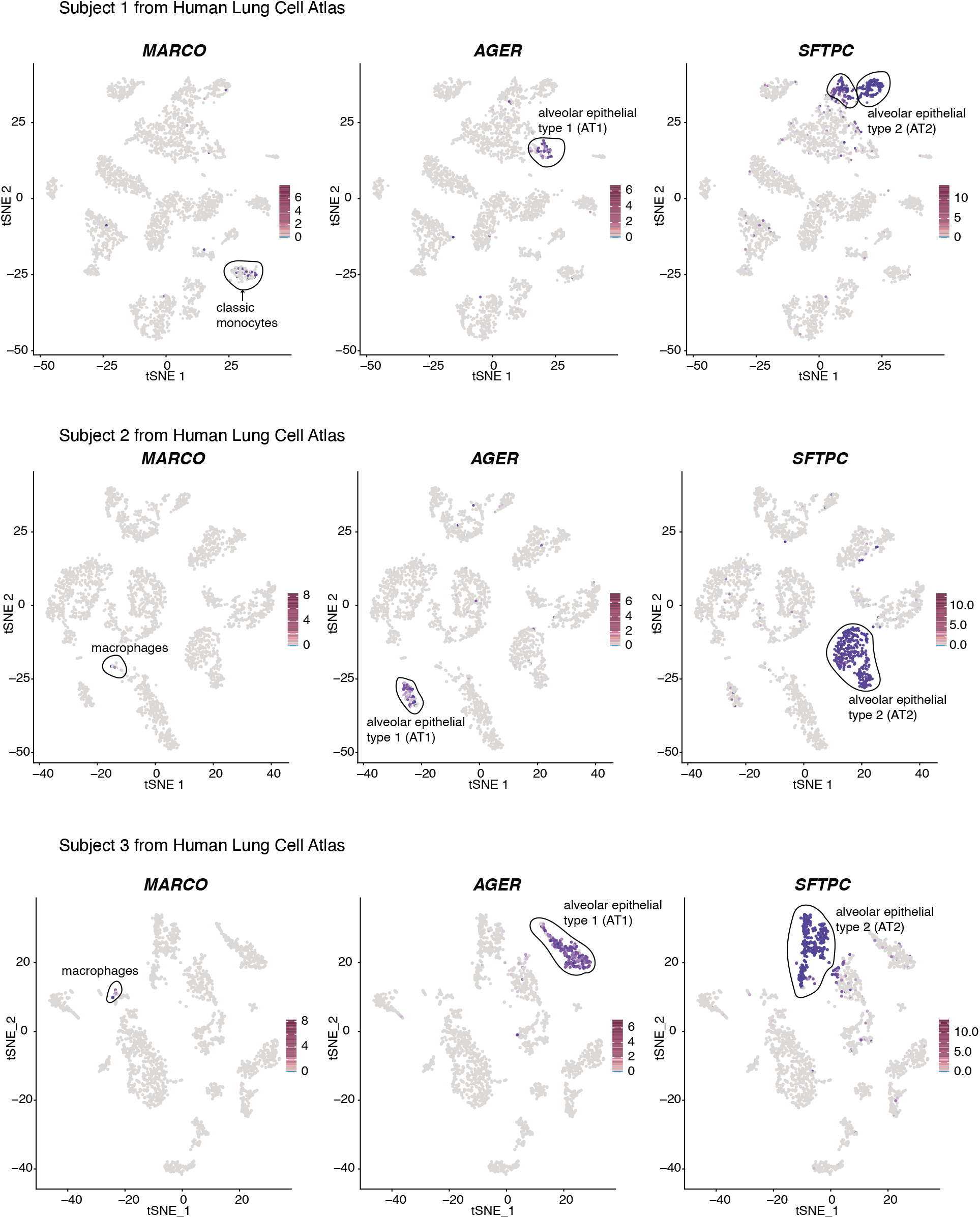
tSNE plots of the human lung cell atlas single-cell RNA-sequencing data across 3 subjects. Each plot is a tSNE projection of all cells in the data set with the color of the points depicting the expression of the gene. Each row of plots is from a different subject. The target cell-type with each marker is labeled on the plots with a circle around the cluster (monocytes/macrophages, AT1 cells, AT2 cells). The three genes identified as cell-type specific markers are *MARCO, AGER*, and *SFTPC*.

## Methods

### Human tissues

Material from autopsies of patients who died of COVID-19 were obtained from family consented research only autopsies performed by the Department of Pathology and Laboratory Medicine at the Hospital of the University of Pennsylvania. De-identified placenta samples were obtained through the Division of Anatomic Pathology at The Children’s Hospital of Philadelphia. Tissues were collected and formalin fixed for 48-72 hours prior to routine processing and paraffin embedding. Tissues were sectioned to 5 um thickness from FFPE blocks to be used in the ISH assays.

### Cell lines and infection

We cultured A549^ACE2^ cells at 37°C and 5% CO2 in RPMI 1640 supplemented with 10% fetal bovine serum (FBS) and 1% penicillin/streptomycin. We cultured Huh7.5 cells at 37°C and 5% CO_2_ in DMEM supplemented with 10% fetal bovine serum, 1% penicillin/streptomycin, and 1% L-Glutamax. For RNA *in situ* experiments, we seeded cells into 2-well chambers (LabTek) at a density of 3,000 cells per well and then infected with SARS-CoV-2 (USA WA1/2020 strain) at an MOI of 1. SARS-CoV-2 was diluted in serum-free RPMI (for A549^ACE^) or DMEM (for Huh7.5) and added to cells for absorption for 1 hour at 37°C. Inoculum was removed and replaced with RPMI with 2% FBS and cells incubated at 37°C. We fixed the cells 24 hours after infection in 4% formaldehyde and PBS for 30 minutes at room temperature. We then washed with PBS two times and permeabilized the samples in 70% ethanol for up to 2 weeks before RNA FISH and RNA FISH HCR. All work with SARS-CoV-2 was performed in a biosafety level 3 laboratory using appropriate personal protective equipment and protocols approved by the Institutional Biosafety Committee and Environmental Health and Safety at the University of Pennsylvania.

### Probe design

We designed RNA FISH HCR probes using RajLab ProbeDesignHD software (code freely available for non-commercial use here: https://github.com/arjunrajlaboratory/ProbeDesign/). This pipeline is implemented in MATLAB and uses a FASTA containing the RNA of interest. The software selects probe sequences according to length and free energy constraints and then excludes probes with complementarity to repetitive elements, human genome, and pseudogenes.

To target the SARS-CoV-2 genome, we referenced the sequence of the first US isolate of SARS-CoV-2 (USA-WA1/2020) from NCBI (Genbank MN985325.1). We used the probe designer described above to design non-overlapping 52-mer oligos with a target Gibbs free energy for binding of −60 (allowable Gibbs free energy [−70, −50]) to the N and ORF1a regions of the SARS-CoV-2 genome, targeting only the 3000-8000 nt region of the latter because it was the most conserved region among the strains circulating at the time as determined using nextstrain^43^. We divided each 52-mer oligo into two non-overlapping 25-mer sequences (removing the middle two nucleotides) and appended split-initiator HCR sequences using a custom matlab script (see Supp. Table 2 for probe sequences). For each probe, we then performed a local blast against the human transcriptome and Nucleic Acids of Coronavirus and other Human Oronasopharynx pathogens (NACHO), a database we created of 562,446 sequences of other viruses that infect the human respiratory tract. All probes in the top 5% of hits based on E-value and bit score were excluded, and the final probe sequences were synthesized from Eurofins at nanomolar-scale. Finally, we resuspended HCR probes to 100 μM in nuclease-free water and then combined these probes into pools each at a final concentration of 2 μM per probe. In the final probe designs, the ORF1a region probe set consisted of 23 probe pairs and the N region probe set consisted of 7 probe pairs.

To target SARS-CoV-2 subgenomic RNAs, we referenced the UCSC Genome Browser for SARS-CoV-2 genome datasets (https://genome.ucsc.edu/covid19.html) and RNA-sequencing datasets^44^ to identify the most frequent junction locations and peri-junction sequences based on the most abundant sgRNA junction spanning reads. We then manually designed 52-mer oligos each spanning a unique leader-body junction for each of the eight canonical subgenomic RNAs generated via discontinuous transcription. We split each 52-mer oligo into two 25-mer sequences and appended split-initiator HCR sequences as outlined earlier (see Supp. Table 2 for probe sequences). The final probes sequences were synthesized and resuspended the same as the SARS-CoV-2 genome-targeting probes.

### Selection of Cell Type Specific Genes

To identify individual cell types from human lung tissue, we reconciled RNA expression level data from multiple single-cell RNA sequencing datasets and identified genes that were both highly expressed and specific to one cell type^36^. Full computational analysis scripts are available here: https://upenn.box.com/s/jtgl47gc869g8w2t0cwc2iov8c520wa0. We then designed probes for specific cell types (see Supp. Table 2) similarly to our SARS-CoV-2 genomic probes and appended split-initiator HCR sequences using our custom matlab script. Final probe sequences were synthesized, resuspended and then combined into pools as outlined earlier.

### HCR RNA FISH

We adapted the previously published HCR v3.0 protocol^24^ for HCR RNA FISH in cultured cells as follows. We used 1.2 pmol each of our pooled HCR RNA FISH probe sets per 0.3 ml of hybridization buffer at 37°C. Our primary hybridization buffer consisted of 30% formamide, 10% dextran sulfate, 9 mM citric acid pH 6.0, 50 μg ml^-1^ of heparin, 1× Denhardt’s solution (Invitrogen) and 0.1% Tween-20. For primary hybridization, we used 300 μl of hybridization buffer containing the appropriate probes per well of a two-well plate (Thermo Fisher Scientific), covered the well with a glass coverslip and incubated the samples in containers humidified with 2× SSC at 37°C overnight (12-16 h). After the primary probe hybridization, we washed samples 4 × 5 min at 37°C with wash buffer containing 30% formamide, 9 mM citric acid pH 6.0, 50 μg ml^-1^ of heparin and 0.1% Tween-20. We then washed the samples 2 × 5 min with 5× SSCT (5× SSC + 0.1% Tween-20) at room temperature and then incubated the samples at room temperature for 30 min in amplification buffer containing 10% dextran sulfate and 0.1% Tween-20. During this incubation, we snap-cooled 0.6 μl per well of individual 3 μM HCR hairpins (Molecular Instruments) conjugated to Alexa Fluor 647 (Alexa647), Alexa Fluor 594 (Alexa594), Alexa Fluor 546 (Alexa546) or Alexa Fluor 488 (Alexa488) in separate PCR tubes by heating at 95°C for 90 s and then either ramp-cooling the sample at a ramp rate of 0.08°C/sec to room temperature in 30 mins or immediately transferring to room temperature for 30 min concealed from light. Next, we pooled the hairpins in 300 μl of amplification buffer to a final concentration of 6 nM each. We added the hairpin solution to samples, placed a glass coverslip on top and then incubated samples at room temperature overnight (12–16 h) concealed from light. After hairpin amplification, we washed samples 5 × 5 min with 5× SSCT, added 100 μl of SlowFade antifade mounting solution containing 50 ng ml^-1^ of DAPI (Invitrogen) with a coverslip and proceeded to imaging the samples.

For HCR RNA FISH on formalin-fixed paraffin-embedded (FFPE) tissues, we obtained tissues fixed via 10% neutral buffered formalin. We deparaffinized tissue sections on slides by first immersing them 2 × 10 min in xylene (Sigma-Aldrich) and then immersing them 2 × 5 min in 100% ethanol. We then transferred the tissues slides to a 3:1 methanol:acetic acid solution at room temperature for 5 mins, washed the slides in nuclease-free water for 3 mins, and then performed antigen retrieval by placing the slides in a solution of 10 mM sodium citrate, pH 6 + 0.1% diethyl pyrocarbonate (Sigma-Aldrich) heated with a 150°C water bath for 15 minutes. After antigen retrieval, we quickly rinsed the slides with 5× SSCT and immediately proceeded to HCR RNA FISH, which we adapted from the previously published HCR v3.0 protocol^24^ for HCR RNA FISH in FFPE tissues as follows. We first pre-hybridized our samples by adding 200 μl of hybridization buffer warmed to 37 °C and incubating at 37 °C for 10 min. While pre-hybridizing, we made our primary hybridization solution containing 0.8 pmol each of our pooled HCR RNA FISH probes per 0.2 ml of hybridization buffer. Our primary hybridization buffer consisted of 30% formamide, 10% dextran sulfate, 9 mM citric acid pH 6.0, 50 μg ml^-1^ of heparin, 1× Denhardt’s solution (Invitrogen) and 0.1% Tween-20. For primary hybridization, we used 50-100 μl hybridization buffer containing the appropriate probes per slide, covered the section with a glass coverslip and incubated the samples in humidified containers at 37 °C overnight (12-16 h). After the primary probe hybridization, we washed samples sequentially in 75% wash buffer (containing 30% formamide, 9 mM citric acid pH 6.0, 50 μg ml^-1^ of heparin and 0.1% Tween-20) + 25% 5× SSCT (5× SSC + 0.1% Tween-20) solution, 50% wash buffer + 50% 5× SSCT solution, 25% wash buffer + 75% 5× SSCT solution and 100% 5× SSCT for 15 min each at 37 °C. We then washed the samples in 5× SSCT at room temperature for 5 min and then incubated the samples at room temperature for 30 min in an amplification buffer containing 10% dextran sulfate and 0.1% Tween-20. During this incubation, we snap-cooled, by heating at 95 °C for 90 s in separate PCR tubes, 0.2 μl per slide of individual 3 μM HCR hairpins (Molecular Instruments) conjugated to Alexa647, Alexa594, Alexa546 or Alexa488 and then either ramp-cooled the sample at a ramp rate of 0.08°C/sec to room temperature in 30 mins or immediately transferred them to room temperature to cool for 30 min concealed from light. After these 30 min, we pooled the hairpins in 100 μl of amplification buffer per slide to a final concentration of 6 nM each. We added the hairpin solution to samples, placed a glass coverslip on top and then incubated samples at room temperature overnight (12–16 h) concealed from light. After hairpin amplification, we washed samples 1 × 5 min in 5× SSCT, 2 × 15 min in 5× SSCT and then 1 × 5 min with 5× SSCT again. We then stained nuclei by adding 200 μl of 5× SSCT containing 50 ng ml^-1^ of DAPI to the samples for 5 mins at room temperature, quenched autofluorescence using the Vector TrueVIEW Autofluorescence Quenching kit according to the manufacturer’s protocol, added a coverslip, and then proceeded to image the samples. We note that the final hairpin concentrations used in these experiments are ten-fold lower than the manufacturer’s protocol, which we optimized to reduce non-specific amplification while still enabling sensitive RNA detection.

### Imaging

We imaged HCR RNA FISH samples on an inverted Nikon Ti2-E microscope equipped with a SOLA SE U-nIR light engine (Lumencor), an ORCA-Flash 4.0 V3 sCMOS camera (Hamamatsu), ×20 Plan-Apo λ (Nikon MRD00205), ×60 Plan-Apo λ (MRD01605) and ×100 Plan-Apo λ (MRD01905) objectives and filter sets for DAPI, Alexa Fluor 488, Alexa Fluor 594 and Atto647N. Our exposure times ranged from 100 ms-200 ms for most of the dyes except for DAPI, for which we used ~50 ms exposures. For experiments in Figs. 2-4, we first acquired tiled images in a single *z*-plane (scan) at ×20 magnification, from which we identified positions containing cells positive for SARS-CoV-2 and returned to those positions to acquire a *z*-stack at ×60 or ×100 magnification. For large area scans, we used Nikon Perfect Focus to maintain focus across the imaging area.

### Image analysis

For quantifying fluorescence intensity in cell culture samples in Fig. 1, we used custom MATLAB scripts available at https://github.com/arjunrajlaboratory/rajlabimagetools. Briefly, our image analysis consisted of manual segmentation of the boundaries for each cell and then quantification of the total fluorescence intensity within that boundary. For plotting in Fig. 1, we normalized the total fluorescence intensity across all pixels in the cell to the total cell area.

For tissue image analysis, we first developed a custom MATLAB pipeline for cropping tiled, single *z*-plane 20×20 scan images taken at ×20 magnification into smaller images. We then used CellProfiler to segment cells using 4’,6-diamidino-2-phenylindole (DAPI) to identify nuclei. We dilated the nuclear objects by a radius of 6 pixels, 7 pixels, and 6 pixels for lung tissue, hilar lymph node tissue and placenta tissue respectively to capture approximately the diameter of one whole cell in the tissue. We measured the position and intensities of the fluorescence signal for each of the SARS-CoV-2 probe sets in each cell. We excluded cells touching image borders. For each cell, we determined the intensity of the SARS-CoV-2 ORF1a probe set by using the cutoff for the upper quartile of pixel intensities across the area of the cell. This processing was necessary because many cells did not have staining throughout the entire cell area. Cellprofiler code is available on the Box for this paper, here: https://upenn.box.com/s/jtgl47gc869g8w2t0cwc2iov8c520wa0.

To determine the fluorescence intensity threshold to label a cell as SARS-CoV-2 positive (in Fig. 2 and 3), we normalized the upper quartile intensity measurements of SARS-CoV-2 ORF1a staining (Alexa647 channel) in each cell to the median intensity across all cells. We then log-transformed the median-normalized data and used the MClust function in R to fit a two-state lognormal mixture model with unequal variance. We evaluated the model fit by an *F* test and selected an appropriate intensity gate that captured only the positive cells in the leftmost distribution. In exposure-matched negative control samples, we found that our *F* test returned insignificant p values (p > 0.05), indicating the presence of only one population of cells with the baseline of background staining. Our intensity gates did not capture any cells in our negative control samples.

For analyses in which we used our SARS-CoV-2 ORF1a probe set with Alexa488 fluorescent hairpins (Fig. 3), we needed a way to exclude autofluorescent background in that channel from our analysis. To do this, we acquired images with a different filter set for which we did not have any dye in the experiment (Alexa594). We then analyzed these images to identify cells with high levels of non-specific signal in this wavelength using the same approach as described for the SARS-CoV-2 ORF1a analysis above. We set a threshold intensity for which cells above had high non-specific signals and removed these cells from our analysis. After removing these cells from the data set, we proceeded with analyzing the ORF1a GFP signal using the analysis pipeline as for Alexa647 described above.

For infected cell type identification analyses shown in Fig. 3, we cropped the large image scans down to individual tiles consisting of roughly one field of view and then segmented the cells as described above using Cell Profiler. We dilated the nuclear segments by a radius of 6 pixels to capture the entire area of each cell. Within each cell, we used the “enhance features” module in CellProfiler to enhance the signal (Alexa647 and Alexa546 channels) from the single-molecule HCR probes for cell type specific genes. We then set a threshold for calling individual HCR spots and assigned the number of spots to each cell. We determined the cell type by plotting the distribution of spot counts for each cell type marker and selecting a threshold that captured the tails of the distributions and adjusted these thresholds manually by referencing HCR RNA FISH images to ensure that our thresholds were reasonably accurate. The threshold for *AGER* was 7 spots (to identify a cell as AT1) and the threshold for *SFTPC* was 6 (to identify a cell as an AT2 cell). Cells that did not meet thresholds or could not be classified based on our parameters were assigned as undetermined.

## Data and code availability

All data and remaining code for these analyses can be found at https://upenn.box.com/s/jtgl47gc869g8w2t0cwc2iov8c520wa0 and upon reasonable request to the corresponding author. All analyses were done in R, MATLAB, or CellProfiler.

## Acknowledgements

We would like to thank Joseph DiRienzi and Igor Tsimberg for their effort obtaining tissues from the research only COVID-19 autopsies and Amy Ziober and Danielle Sharpe for providing slides for the ISH experiments. We would also like to acknowledge the TTAB staff that helped to collect the tissues including Federico Valdivieso, Lauren Schmucker, Philip Feldman, Alexa Craig, Jenny Chaiyasid, and Nina Thakur. SMS acknowledges support from the NIH Director’s Early Independence Award DP5OD028144. SMS and SC acknowledge support from the Collaborative Research Grant from the University of Pennsylvania’s Institute for Regenerative Medicine and the Dean’s Innovation Fund. BE acknowledges support from NIH F30 CA236129, NIH T32 GM007170, and NIH T32 HG000046. SRW acknowledges support from R01 AI140442. SC acknowledges support from the NIH and Mark Foundation (19-011-MIA); Dean’s Innovation Fund; Linda and Laddy Montague; BWF; NIAID (5R01AI140539, 1R01AI1502461, and R01AI152362); NCATS, the Fast Grants Award from Mercatus; and the Bill and Melinda Gates Foundation for funding. S.C. is an investigator in the Pathogenesis of Infectious Diseases from the Burroughs Wellcome Fund.

## Author Contributions

KKA, BLE, JL, and SMS conceived and designed the project with valuable input from DLS, RLL, SRW, KTM, SC, and SMS. KKA, BLE, DLS, and JK performed the RNA FISH HCR experiments in tissues. CEC, SC, and SRW performed and supervised cell culture infection experiments with SARS-CoV-2. SR performed the single-molecule RNA FISH experiments. EKS performed analysis of the lung atlas single-cell RNA-sequencing data. NAK, LAL, RLL, MF, and KTM led the clinical protocols and procurement of tissues for this research. KKA and DLS constructed the image analysis pipeline and implemented it on the data. KKA and SMS wrote the manuscript.

